# CC-seq: Nucleotide-resolution mapping of Spo11 DNA double-strand breaks in *S. cerevisiae* cells

**DOI:** 10.1101/2023.10.31.564935

**Authors:** George G. B. Brown, William H. Gittens, Rachal M. Allison, Antony W. Oliver, Matthew J Neale

## Abstract

During meiosis Spo11 generates DNA double-strand breaks to induce recombination, becoming covalently attached to the 5′ ends on both sides of the break during this process. Such Spo11 “covalent complexes” are transient in wild-type cells, but accumulate in nuclease mutants unable to initiate repair. The CC-seq method presented here details how to map the location of these Spo11 complexes genome-wide with strand-specific nucleotide-resolution accuracy in synchronised *Saccharomyces cerevisiae* meiotic cells.

## 1. Introduction

During meiosis, homologous recombination (HR) between non-sister chromatids provides a mechanism for generating genetic diversity. HR is initiated deliberately, yet in a non-random manner, via the generation of DNA double strand breaks (DSBs) created by the evolutionarily conserved Spo11 enzyme. Failure to initiate or properly repair Spo11 DSBs leads to chromosomal mis-segregation in meiosis creating aneuploidy. In humans, aneuploidy is a cause of genetic disorders such as Down’s syndrome.

Spo11 is part of the topoisomerase family of proteins that create DNA breaks via the formation of a 5’-phosphotyrosyl-linked covalent complex (CC) ***(1)***. The Spo11-linked DNA molecule is then endonucleolytically removed via the Mre11 complex (Mre11, Rad50, Xrs2)—an activity dependent upon the Sae2 protein ***(2–4)***, the orthologue of human CtIP ***(5)***. Nicks created by Mre11 flanking the Spo11 DSB act as initiation sites for bidirectional resection by Mre11 and Exo1 in the 3’ to 5’ and 5’ to 3’ directions, respectively ***(2***; ***6)***. The ssDNA that is revealed acts as a substrate for loading RecA-family recombinases such as Dmc1 and Rad51, which promote Spo11-DSB repair by catalysing strand invasion between homologous chromosomes, leading to crossover and non-crossover repair outcomes.

Recombination promotes genetic variation and accurate meiotic chromosome segregation. Thus, understanding the mechanisms that govern spatial and temporal patterns of Spo11-DSB formation provide insight into patterns of genetic change arising within sexually breeding populations and the potential causes of genetic instability and aneuploidy.

Since the discovery of Spo11, several complementary methods for assessing genome-wide patterns of Spo11 DSBs have been developed—some of which have been applied to numerous experimental systems. These include:

⍰ DNA microarray analysis of the accumulation of Spo11-bound DNA in the nuclease-deficient *rad50S* background in *S. cerevisiae* ***(7)***.
⍰ Single-stranded DNA (ssDNA) microarray mapping using RPA ChIP-chip ***(8)*** or ssDNA enrichment ***(9)*** in strand-invasion deficient *S. cerevisiae* backgrounds (*dmc1*∆ and/or *rad51*∆), and related sequencing techniques in *Mus musculus* ***(10)***.
⍰ High-resolution Spo11-oligo mapping in, *S. cerevisiae* ***(11–13)***, *Schizosaccharomyces. pombe* ***(14)***, *M. musculus* ***(15)***, and *Arabidopsis thaliana* ***(16)***.
⍰ High-resolution Spo11-DSB mapping methods: S1-Seq in *M. musculus* ***(17)*** *and S. cerevisiae* ***(18***; ***19)*** and End-Seq in *M. musculus* ***(20)***.

Here, we describe a detailed protocol for CC-seq ***(21***; ***22)***, an antibody-free method for determining the genome-wide locations of Spo11-DSB formation in meiotic cells with single-nucleotide accuracy (Figure 1). The method exploits the electrostatic interaction between protein and silica to enrich Spo11-CC linked double-stranded DNA molecules that accumulate when the Mre11-dependent nucleolytic repair pathway is prevented (i.e. in *sae2*∆, *rad50S*, or *mre11* nuclease-dead cells). Pilot data indicate that CC-seq, whilst optimised for analysis of Spo11-DSBs in *S. cerevisiae* meiotic cells, is also transferable to other single-cell eukaryotes such as *S. pombe* and *Lachancea kluyveri*.

**Figure 1.**
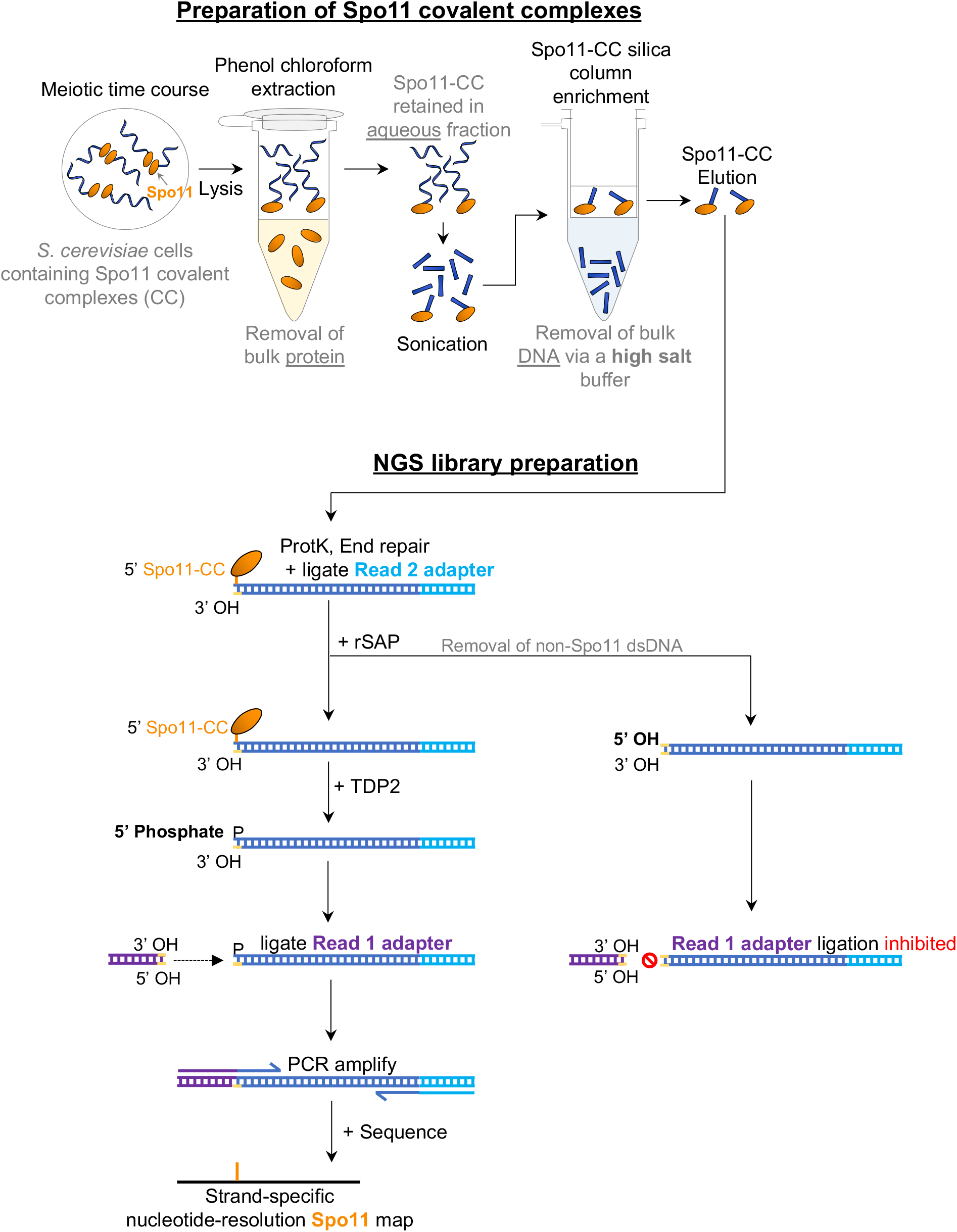
Overview of the CC-seq methodology. Top, Cartoon showing the general steps of Spo11-CC enrichment via cell lysis, phenol chloroform extraction, sonication and enrichment on silica membrane. Bottom, Cartoon showing the sequential DNA library preparation stages (see text for details). Treatment with rSAP prevents adapter ligation to non-Spo11 ends, increasing specificity.

Briefly, DNA containing protein-DNA covalent complexes (CC-DNA) from *S. cerevisiae* meiotic cells is prepared using an SDS and phenol chloroform extraction, fragmented by sonication, then passed through a silica membrane in the presence of a high-salt buffer similar to prior work ***(1***; ***7)***. Following a series of high-salt washes to remove background non-CC-DNA, enriched Spo11-linked DNA molecules are eluted with detergent. Adapters suitable for next-generation sequencing on the Illumina® platform are ligated to both ends of each Spo11-CC dsDNA fragment, utilising the 5’-phosphotyrosyl-unlinking activity of recombinant vertebrate TDP2 ***(23)*** to differentiate sonicated from Spo11-linked DNA ends. Finally, resulting sequence reads are aligned to the *S. cerevisiae* reference genome generating a strand-specific nucleotide-resolution map of where Spo11-DSBs form within the genome (Figure 2) (see note 1).

**Figure 2.**
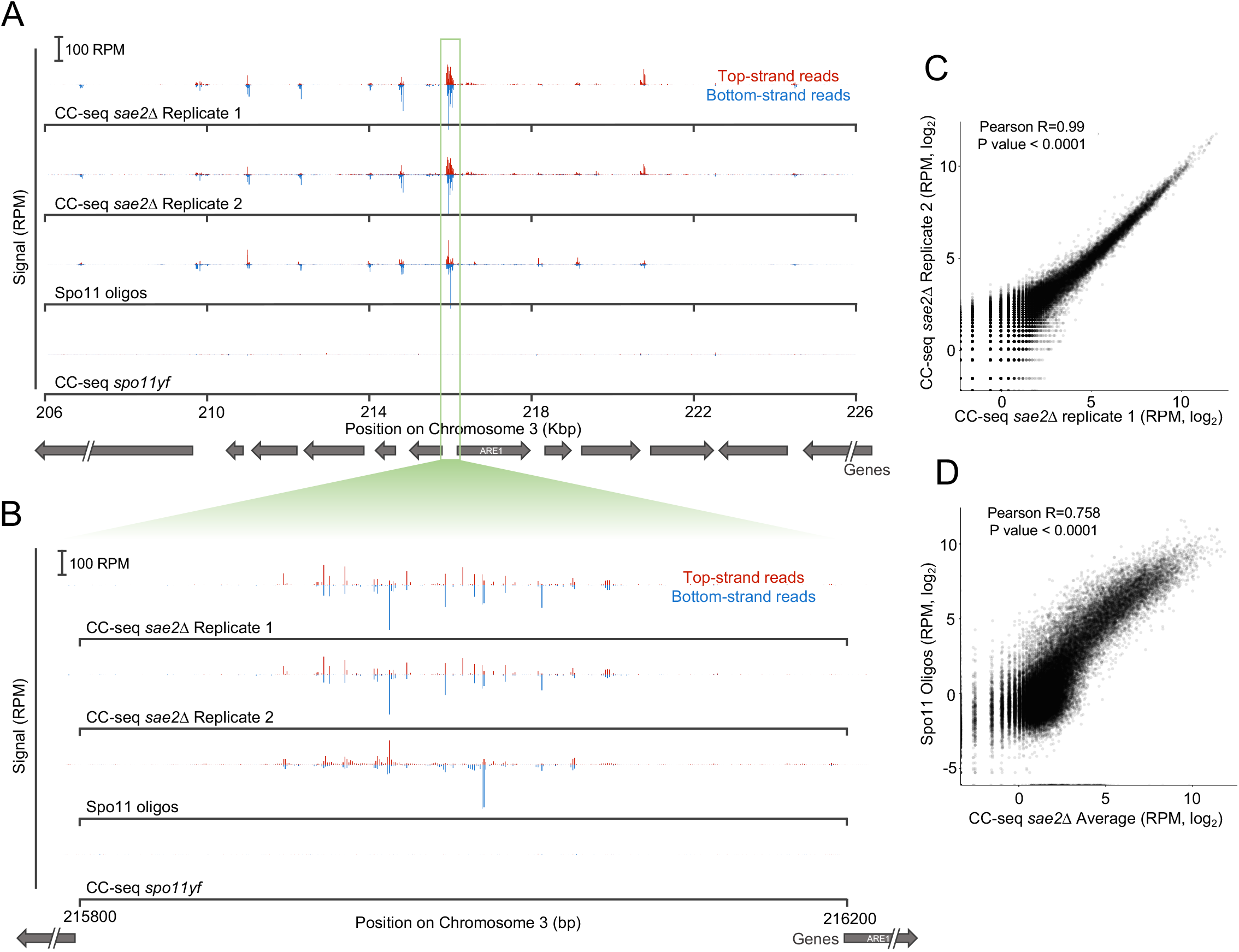
Reproducibility of CC-seq maps. A-B, Spatial reproducibility of CC-seq maps compared to Spo11-oligos (GSE84696; ***(13)***). Mapped reads were normalised to reads per million (RPM). The bottom strand reads were offset by -1 bp. (A) 20 Kbp region of chromosome III is shown as an example of the visual reproducibility between two independent biological repeats of the *sae2*∆ datasets. B, Nucleotide reproducibility of CC-seq reads at the *ARE1* hotspot (a representative 400 bp region of chromosome III). C-D, Normalised, mapped reads were summed in 250 bp non-overlapping windows. (C), Quantitative comparison of CC-seq maps for the same two *sae2*∆ biological replicates (Pearson *r* = 0.99). (D), Quantitative comparison between the average of two *sae2*∆ biological replicates and Spo11-oligos (GSE84696; ***(13)***) (Pearson *r* = 0.758).

### 2. Materials

### 2.1.1. General

1. General lab equipment, including: shaking incubator, temperature controlled floor standing centrifuge and rotors, swinging bucket bench top centrifuge, vacuum aspirator, dry heat block, shaking heat block, water bath, 5 mL microcentrifuge with 5 mL tubes with suitable rotor, mini-PCR tube centrifuge, PCR tube thermocycler, 200 μL PCR tubes
2. Qubit® 2 (Invitrogen #Q32866)
3. Qubit® dsDNA HS Assay Kit (Invitrogen #Q32854)
4. Qubit® Assay Tubes (Invitrogen #Q32856)
5. 1 × TE: 10 mM Tris, 1 mM EDTA pH 7.8

### 2.2. Preparation of Spo11 covalent complexes

#### 2.2.1. Meiotic time course and cell collection

1. Yeast Growth media: YPD: 1% Yeast extract, 2% Peptone, 2% D-glucose BYTA: 1% Yeast extract, 2% tryptone, 2% Potassium (KAc), 50 mM Potassium phthalate SPM: 0.3% Potassium (KAc), Raffinose 0.02%, Adenine 5 μg/mL, Arginine 5 μg/mL, Histidine 5 μg/mL, Leucine 15 μg/mL, Tryptophan 5 μg/mL, Uracil μg/mL
2. Copper(II) sulphate (Merck #C1297)

#### 2.2.2. Cell spheroplasting and fixing

1. Zymolyase 100T (Amsbio #120493-1) dissolved at 50 mg/mL in 35% glycerol, 50 mM NaHPO4 buffer pH 7.2, 0.5 M sucrose
2. Beta mercapto-ethanol (βME) (Merck #M6250)
3. Spheroplasting buffer: 1 M sorbitol, 50 mM NaHPO4 pH 7.2, 10 mM EDTA

#### 2.2.3. SDS cell lysis

1. STE: 2% SDS, 0.5 M Tris.HCl, pH 8.1, 10 mM EDTA, Bromophenol blue 20 μg/mL
2. Pestle for 1.5 mL tubes (VWR #431-0094)

#### 2.2.4. Phenol chloroform DNA extraction

1. Phenol; Chloroform; lsoamyl alcohol in the ratio of 25:24:1 (v/v/v); saturated with 100 mM Tris pH 8.0; contains ∼0.1% 8-hydroxyquinoline (Merck #77617)

#### 2.2.5. DNA sonication and DNA quantification

1. Covaris M220 machine (Covaris #500295)
2. 1 mL milliTUBE 1 mL AFA Fiber (Covaris #520135)
3. XT milliTUBE 1 mL holder (Covaris #500348)
4. 200 μL PCR tube
5. RNase A pH 7.4 10 mg/μL (Merck #R4875), in 9 mM sodium acetate buffer pH 5.2, 100 mM Tris-HCl pH 7.4. Heat treated to 100°C for 15 min, then slowly cooled

#### 2.2.6. Spike in preparation

1. Lambda DNA (∼500 μg/mL) (NEB #N3011)
2. Human hTERT RPE-1 cells (ATCC #CRL-4000)
3. Dulbecco’s Modified Eagle’s Medium DMEM/F-12 (Merck #D6421-500 mL)
4. 10% Foetal Calf Serum (FCS)
5. Penicillin-Streptomycin (10,000 U/mL) (Gibco™ #15140122)
6. Etoposide (Merck #E1383-100MG)
7. Syringe and needle (21 gauge)

#### 2.2.7. Spo11 covalent complex enrichment

1. 10 × column binding buffer: 3 M NaCl,1% Sarkosyl, 2% TritonX100
2. TEN: 10 mM Tris HCl pH 7.8, 1 mM EDTA, 0.3 M NaCl
3. TES: 10 mM Tris HCl pH 7.8, 1 mM EDTA, 0.5% SDS

#### 2.2.8. Deproteinisation of Spo11 covalent complexes

1. Proteinase K (FisherScientific #10407583) dissolved at 50 mg/mL in 50% glycerol, 20 mM Tris-HCl pH 7.8
2. Glycogen 20mg/μL (NEB #R0561)
3. 3 M NaOAc

### 2.3. NGS library preparation

1. 10 mM Tris pH 8.2
2. Ultra II DNA Library Prep Kit for Illumina® (NEB #E7645)
3. AMPure® XP magnetic beads (Beckman Coulter #A63882)
4. PCR tubes (200 μL)
5. DynaMag block (200 μL) (ThermoFisher #492025)
6. rSAP and rCutSmart buffer (NEB #M0371)
7. Adapter dilution buffer: 10 mM NaCl, 10 mM Tris HCl pH 8.2
8. 2 × Tdp2 buffer: 100 mM NaOAc, 100 mM TrisOAc, 2 mM MgOAc, 2 mM DTT (FisherScientific #10592945), 200 μg/mL Recombinant albumin (NEB #B9200S).
9. NEBNext® Multiplex Oligos for Illumina® (Index Primer Set 1) (NEB #7335S/L)
10. Bioanalyzer instrument (Agilent #G2939BA), Bioanalyzer Chip Priming Station (Agilent #5065-4401), 2100 Expert SW Laptop Bundle (Agilent #G2953CA), Agilent High Sensitivity DNA Kit (Agilent #5067-4626)

### 2.4. NGS data processing

1. terminalMapper (https://github.com/Neale-Lab)
2. CCTools (https://github.com/Neale-Lab)

## 3. Method

All centrifugation, incubation and handling steps have been performed at room temperature (∼22°C) unless stated otherwise.

### 3.1. Preparation of Spike-in DNA, custom adaptors and recombinant TDP2

#### 3.1.1. Spike-in DNA preparation (Optional)

To estimate relative Spo11-DSB yield between libraries, etoposide (VP16)-treated human DNA, used as a spike-in, is prepared as follows:

1. Plate hTERT RPE-1 cells (or other adherent human cells) on 4 × 15 cm plates at a density of 0.6 × 10^6^ cells per plate and incubated at 37°C in 30 mL DMEM/F12 supplemented with 10% FCS and Pen/Strep (complete media) for 72 hours.
2. Cell confluency should be 60-70%, with ∼5 × 106 cells per plate (20 x 106 cells total).
3. Treat each plate for 30 min with a final concentration of 100 μM etoposide in 20 mL complete media.
4. Remove treatment media and immediately add 10 mL ice-cold 70% ethanol-PBS.
5. Immediately scrape cells into this 70% ethanol-PBS buffer.
6. Pool cells from all 4 plates in a 50 mL falcon tube on ice.
7. Centrifuge for 5 min at 400 × *g* at 4°C.
8. Aspirate supernatant and resuspend pellet in 4 mL ice-cold 70% ethanol-PBS. Transfer to a 5 mL eppendorf (or 10 mL falcon).
9. Centrifuge for 5 min at 400 × *g* at 4°C.
10. Aspirate supernatant.
11. Process ethanol-fixed cells according to “SDS cell lysis” and “Phenol-chloroform extraction of DNA” steps above.
12. Resuspend DNA by incubating in a thermomixer (5 min 1000 RPM, 45°C).
13. Pipette suspended DNA into a 1 mL milliTUBE AFA Fiber.
14. Sonicate using a Covaris M220 and XT milliTUBE 1 mL holder using the following settings (see note 2):
  - Target BP range: 200-700 bp
  - Duty Cycle: 10%
  - Intensity/Peak power incidence: 75W
  - Cycles/burst: 200
  - Time: 10 min
  - Bath Temp: 7°C
15. Pipette 12 μL sonicated VP16 treated human DNA into a PCR tube.
16. Add 3 μL RNase A (3.3 mg/mL) to a final concentration of 0.66 mg/mL.
17. Incubate at 37°C for 20 minutes.
18. Measure the RNase-treated human DNA sample to calculate the DNA concentration via Qubit® using the HS dye protocol (see note 3).
19. Using the Qubit® DNA concentration readings, dilute the RNase-treated human DNA to 80 μg/mL in 1 × TE pH 7.6 (see note 4).

To estimate noise differences between libraries, Lambda DNA is prepared as follows:

1. Mix 2.5 mL of 500 μg/mL Lambda DNA with 2.5 mL 1 × TE pH 7.6 in a 5 mL tube, and vortex for 20 seconds.
2. Use a syringe to pipette up and down 40x to help homogenise the Lambda DNA.
3. Aliquot Lambda DNA into 5 x 1 mL milliTUBE AFA Fiber.
4. Sonicate using a Covaris M220 and XT milliTUBE 1 mL holder using the following settings (see note 2):
  - Target BP range: 200-700 bp
  - Duty Cycle: 10%
  - Intensity/Peak power incidence: 75W
  - Cycles/burst: 200
  - Time: 10 min
  - Bath Temp: 7°C
5. Repeat the previous step for each Lambda DNA 1 mL aliquot.
6. Pool aliquots (5 mL total) in a 5 mL microcentrifuge tube, and vortex heavily.
7. Pipette 2.5 μL of Lambda DNA and mix with 7.5 μL DD water in a PCR tube.
8. Measure the diluted Lambda DNA concentration after sonication via Qubit® using the HS dye protocol.
9. Using the Qubit® DNA concentration readings, dilute the Lambda DNA to 150 μg/mL in 1 × TE pH 7.6.

In order to estimate Spo11-DSB signal: noise between libraries, etoposide (VP16)-treated human DNA (signal) is mixed with Lambda DNA(noise) at a 1:50 ratio prior to “Spo11 covalent complex enrichment”, prepared as follows:

1. Mix 187.5 μL of VP16 treated Human hTERT RPE-1 DNA (80 μg/mL) with 5 mL Lambda DNA (150 μg/mL) (see note 5).
2. Prepare in advance and store 200 μL aliquots at –80°C.

#### 3.1.2. Custom Read 1 and Read 2 adapter generation

1. Custom oligonucleotides were generated using https://www.idtdna.com dissolved at a final concentration of 100 μM in 1 × TE pH 7.6:
  - Read1_Adapter_P5_top: ACACTCTTTCCCTACACGACGCTCTTCCGATCT
  - Read1_Adapter_5’-dephospho-P5_bottom:
  - GATCGGAAGAGCGTCGTGTAGGGAAAGAGTGT/3InvdT/
  - Read2_Adapter_P7_top:
  - /5Phos/GATCGGAAGAGCACACGTCTGAACTCCAGTCAC/3InvdT/
  - Read2_Adapter_P7_bottom: GTGACTGGAGTTCAGACGTGTGCTCTTCCGATCT
2. For Read 1 and Read 2 adapter generation mix 6 μL of top and bottom oligonucleotides.
3. Add 26 μL of 1 × TE pH 7.6.
4. Add 2 μL 1 M NaCl and mix thoroughly.
5. Anneal oligonucleotides by heating for 5 minutes at 100°C then leave to cool slowly for 2 hours.
6. Final adapter concentration: 15 μM in 50 mM NaCl 1 × TE pH 7.6.
7. Prepare in advance and store 40 μL aliquots at –20°C.

#### 3.1.3. 14M_zTDP2CAT expression and purification

1. To generate 14M_zTDP2CAT(240 μM) a codon-optimised synthetic gene encoding the catalytic domain of zebrafish TDP2 (amino acids 120-369)—where 14 individual single point mutations have been introduced in order to ‘humanise’ the encoded enzyme ***(24)***—was purchased from GeneArt (ThermoFisher Scientific, UK).
2. The coding sequence was sub-cloned into the NdeI and EcoR1 sites of pAWO-SUMO-3C, an in-house modified version of pET-17b that encodes an in-frame, human rhinovirus 3C-protease (HRV-3C) cleavable N-terminal fusion with yeast Smt3 prepended with a His6 affinity tag. The plasmid is available from AddGene: https://www.addgene.org/200512/
3. Expression and purification of the recombinant protein followed the protocol described for m2hTDP2CAT ***(25)***, but replacing SENP1 protease with HRV-3C.
4. To generate 30 μM aliquots dilute 120 μL of 14M_zTDP2CAT(240 μM) into 840 μL x2 Tdp2 buffer containing 50% glycerol.
5. Prepare in advance and store 100 μL aliquots at –80°C.

### 3.2. Preparation of Spo11 covalent complexes

#### 3.2.1. Meiotic time course and cell collection

1. From a –80°C glycerol stock, streak out yeast cells onto a YPD plate. Incubate for 2 days at 30°C.
2. In a glass culture tube, inoculate 4 mL YPD liquid with a single colony taken from the YPD plate. Incubate 24 h at 30°C with shaking (250 RPM).
3. Dilute the cell culture 1 in 50 with water to measure cell density (OD_600_). After 24 h cells usually reach an OD_600_ of ∼20.
4. In a 2 L flask, inoculate 150 mL BYTA to a cell density (OD_600_) of 0.3 by adding 45 OD_600_ units (∼2.25 mL) of cell culture from the YPD overnight. Incubate 15 h at 30°C with shaking (250 RPM).
5. Dilute pre-sporulation BYTA culture 1 in 20 with water to measure cell density (OD_600_). After 15 h cells usually reach an OD_600_ of ∼6.5.
6. Collect 625 OD_600_ units (∼100 mL) of pre-sporulation BYTA culture by centrifugation (5 min at 3250 × *g*) in a 250 mL centrifuge bottle and drain supernatant (see note 6).
7. Add 200 mL water, seal bottle and shake vigorously to resuspend cells.
8. Pellet cells again (5 min at 3250 × *g*), drain, and resuspend in 250 mL SPM at 30°C (final OD_600_ of ∼2.5) by shaking vigorously.
9. Pour cell suspension into a new 2 L flask and incubate at 30°C with shaking (250 RPM).
10. After 2 hours add 50 μL of 250 mM Copper (II) sulphate (50 μM final concentration) to induce meiosis via the *P*_*CUP1*_*IME1* promoter.
11. Monitor DNA replication timing by sampling cells at 0, 1, 1.5, 2 and 4 hours after adding Copper (II) sulphate by pipetting 400 μL of culture into 1 mL of 100% EtOH and processing for standard *S. cerevisiae* FACS analysis protocol ***(26)***.
12. Collect 50 mL cells at desired time points (usually between 2–4 hours after adding Copper (II) sulphate) by pouring into a 50 mL falcon tube and incubate in ice-water for 5 minutes.
13. Centrifuge (5 min at 3250 × *g*, 4°C) to pellet the cells. Pour off supernatant.
14. Store cell pellet in a –20°C freezer for at least 8 hours.

#### 3.2.2. Cell spheroplasting and fixing

From this point the protocol describes a single sample. In practice 4-8 samples are routinely processed in parallel.

1. Pre-aliquot 3.75 mL 100% EtOH into a 5 mL Eppendorf tube, and place in the freezer (-20°C).
2. Mix 1.2 mL spheroplast buffer, 6 μL zymo100T, 12 μL βME to a final concentration of zymo100T 250 μg/mL and βME 1% just before use.
3. Add 1.2 mL of Spheroplast solution to the frozen cell pellet.
4. Resuspend the cell pellet by vortexing.
5. Incubate the suspended cells in a water bath at 37°C for 30 min.
6. Fix the cells by pipetting the cell suspension into the pre-aliquoted 3.75 mL EtOH (-20°C). Shake vigorously.
7. Incubate in the freezer (-20°C) for 10 minutes.
8. Collect cells by centrifugation (1 min at 1000 × *g*). Aspirate supernatant.

#### 3.2.3. SDS cell lysis

1. Add 800 μL STE.
2. Completely resuspend into a thick paste using a pestle.
3. Add 1600 μL STE and mix thoroughly by inversion.
4. Incubate in a dry heat block at 65°C for 10 min, invert once during incubation.
5. Cool on ice for 1 min, then place at room temperature.

#### 3.2.4. Phenol chloroform DNA extraction

1. Add 1250 μL phenol; chloroform; isoamyl alcohol. Cap tube securely and shake vigorously for 10 seconds.
2. Incubate cell lysate at room temperature for 5 minutes.
3. Shake vigorously for 10 seconds.
4. Separate the cell lysate into the aqueous, interphase and phenol phase by centrifugation (15 min at 21000 × *g*)
5. Carefully remove 2000 μL of supernatant, using a P1000 tip with ∼5 mm cut off the tip to increase the bore diameter, and pipette into a clean 5 mL tube containing 3.3 mL 100% EtOH.
6. Mix by inversion 20x. A large stringy DNA precipitate will form.
7. Collect DNA by gentle centrifugation (1 min at 21000 × *g*), aspirate supernatant.
8. Wash the DNA pellet with 5 mL 70% EtOH carefully.
9. Collect DNA by centrifugation (1 min at 21000 × *g*), aspirate supernatant.
10. Dry at room temperature for 10 min.
11. Add 1 mL 1 × TE (pH 7.8) to the DNA pellet and leave at 4°C to soak overnight.
12. (Optional) Break point (see note 7).

#### 3.2.5. DNA sonication and DNA quantification

1. Resuspend DNA by incubating in a thermomixer (5 min 1000 RPM, 45°C).
2. Pipette suspended DNA into a 1 mL milliTUBE AFA Fiber.
3. Sonicate using a Covaris M220 and XT milliTUBE 1 mL holder using the following settings (see note 2):
  - Target BP range: 200-700 bp
  - Duty Cycle: 10%
  - Intensity/Peak power incidence: 75W
  - Cycles/burst: 200
  - Time: 15 min
  - Bath Temp: 7°C
4. Pipette 12 μL sonicated DNA into a PCR tube
5. Add 3 μL RNase A (3.3 mg/mL) to a final concentration of 0.66mg/mL.
6. Incubate at 37°C for 20 minutes.
7. Analyse 10 μL of RNase-treated DNA using a 1% agarose gel with ethidium bromide concentration at 0.5 μg/mL to determine sonication efficiency (Figure 3) (see note 2).
8. Measure the RNase-treated DNA sample to calculate the DNA concentration via Qubit® using the HS dye protocol (see note 8).

**Figure 3.**
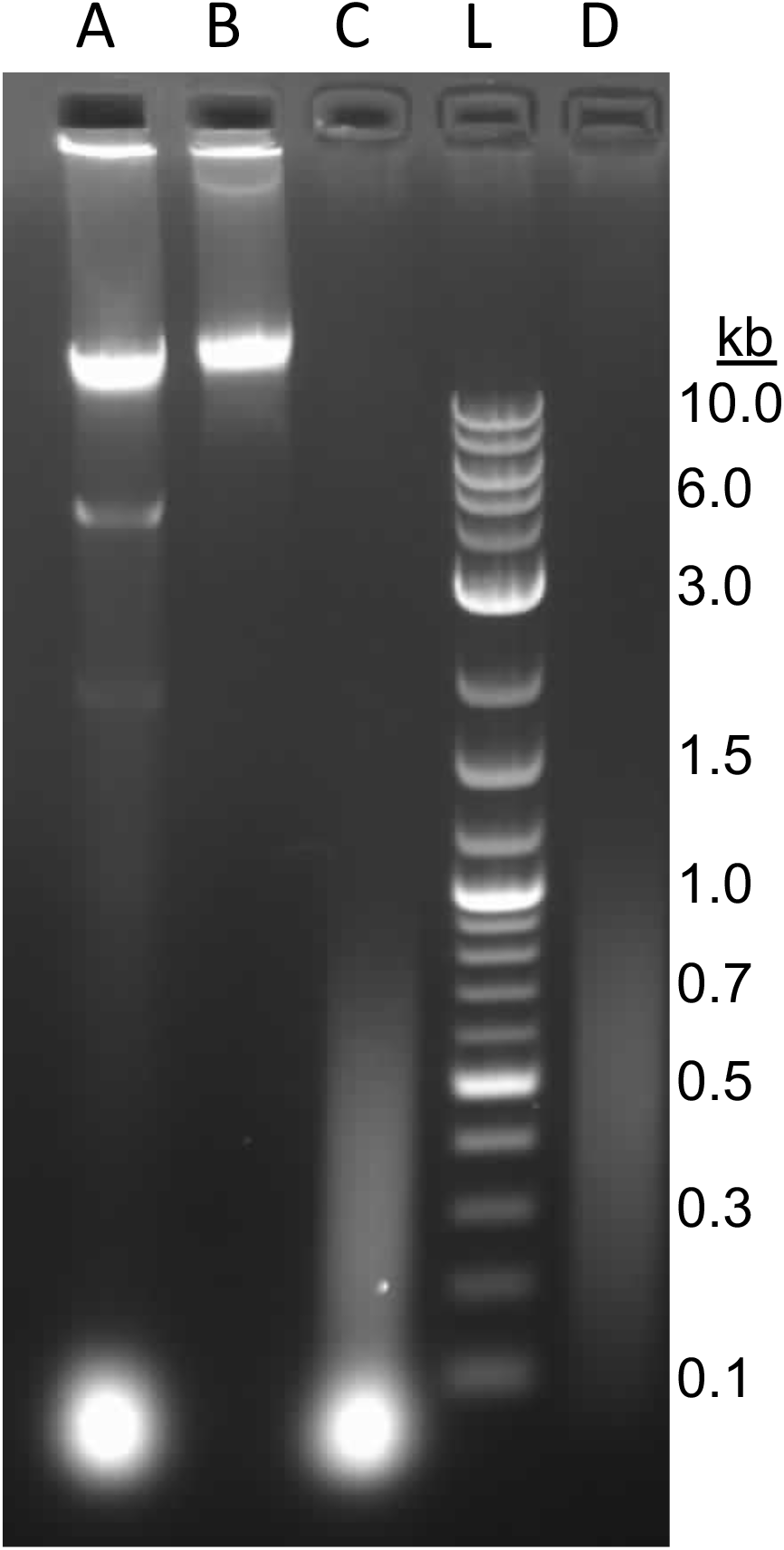
DNA size distribution for a typical CC-seq library. Samples are separated on a 1% agarose gel containing 0.5 μg/mL EtBr for 1 hour at 100 V. Library samples were treated as follows: Lane A, untreated. Lane B, RNase treated only. Lane C, sonicated only. Lane L, 1 μg of log2 DNA ladder (NEB #N3200). Lane D, RNase treated and sonicated.

#### 3.2.6. Spo11 covalent complex enrichment

1. Pipette 900 μL of sonicated DNA into a 1.5 mL microcentrifuge tube.
2. Add 110 μL of the column binding buffer.
3. (Optional) add a DNA spike control to a final % DNA concentration of 0.1% VP16-treated human DNA and 5% Lambda DNA (see section 3.1.1).
4. Top up the column binding buffer and sonicated DNA solution with 1 × TE pH 7.6 to 1.1 mL and mix thoroughly.
5. Load two Qiaprep silica columns each with 550 μL of sonicated DNA solution and incubate for 5 minutes.
6. Centrifuge (1 min at 21000 × *g*).
7. Reload each column by pipetting the flow-through back onto the same column and incubate for 5 minutes.
8. Centrifuge (1 min at 21000 × *g*).
9. Wash column with six sequential washes with 600 μL TEN. Centrifuge (1 min at 21000 × *g*) each time and discard flowthrough.
10. Perform a final dry centrifugation (1 min at 21000 × *g*) to remove any residual wash buffer.
11. Add 125 μL TES (warmed to 60°C) to column and incubate at room temperature for 5 min to elute Spo11-CC DNA.
12. Centrifuge (1 min at 21000 × *g*).
13. Repeat steps 11-12 with a second aliquot of 125 μL TES.
14. Pool eluates (500 μL total) in a 2 mL microcentrifuge tube.

#### 3.2.7. Deproteinisation of Spo11 covalent complexes

1. Add 10 μL Proteinase K (50 mg/mL) to a final concentration of ∼1 mg/mL.
2. Incubate in a thermomixer (1 hour, 800 RPM, 50°C).
3. Allow samples to cool to room temperature
4. Add 5 μL of glycogen (20 mg/mL) to a final concentration of ∼0.2 μg/μL and vortex.
5. Add 50 μL of 3 M NaOAc (265.5 mM final) and vortex.
6. Add 1.41 mL of 100% EtOH and vortex.
7. Incubate at -20°C overnight to precipitate the DNA. Caution: For steps 8 and 10 the DNA pellet is translucent and difficult to see so proceed with care.
8. Collect precipitated DNA via centrifugation (60 min at 21000 × *g*, 4°C), aspirate supernatant.
9. Wash the DNA with 70% EtOH (-20°C) slowly, do not vortex.
10. Collect precipitated DNA via centrifugation (15 min at 21000 × *g*, 4°C), aspirate supernatant.
11. Air dry at room temperature for 10 minutes.
12. Add 25 μL 10 mM Tris HCl pH 8.2.
13. Resuspend DNA by thermomixing (5 min 1000 RPM, 45°C).
14. Remove 25 μL to a new PCR tube.
15. (Optional) Measure the DNA concentration via Qubit® using the HS dye protocol (see note 9).
16. (Optional) Break point (see note 7).

### 3.3 NGS library preparation

This protocol describes one library but normally 4-8 libraries can be processed in parallel.

#### 3.3.1. DNA end repair and A tailing

To 25 μL of CC-enriched sample (thawed at room temperature if previously frozen):

1. Add 3.5 μL NEBNext® Ultra II End Prep Reaction Buffer.
2. Add 1.5 μL NEBNext® Ultra II End Prep Reaction.
3. Vortex to mix and quickly spin down in mini-PCR centrifuge.
4. Place in a thermocycler and run the following program: 30 min at 20°C, 30 min at 65°C.

#### 3.3.2. Read 2 adapter ligation

1. During the 60 minute incubation prepare a mastermix of the following components: 15 μL NEBNext® Ultra II Ligation Master, 0.5 μL NEBNext® Ligation Enhancer and keep on ice (see note 11).
2. In a separate tube, mix 1 μL of **Read 2 P7** adapter (15 μM) with 9 μL of adapter dilution buffer and mix thoroughly. This ten-fold diluted stock is sufficient for 8 libraries. Do **not** add adapters to the master mix.
3. Remove sample from thermocycler and cool to room temperature. Add 15.5 μL of the NEBNext® master mix.
4. Add 1.25 μL of diluted Read 2 P7 adapter (1.5 μM).
5. Mix thoroughly by vortexing for 2 seconds and quickly spin down
6. Incubate at 20°C overnight in a thermocycler with the heated lid off.

#### 3.3.3. Generic AMPure® XP bead clean up protocol

Refer to main protocol sections for specific volumes and buffers to use at each purification stage.

1. Take AMPure® XP beads out of the fridge at least 30 mins before use and shake vigorously 30-40 times to completely suspend. Add ***specified volume*** AMPure® XP beads (0.83 × final concentration) and vortex for 2 seconds.
2. Invert the PCR tubes until the beads are completely suspended.
3. Incubate samples on the bench for 5 minutes (see note 10).
4. Place samples on a PCR tube DynaMag block (200 μL size).
5. After 5 minutes pipette off and discard the supernatant being careful not to disturb the beads on the sides of the tube.
6. Add 200 μL of 80% freshly prepared EtOH to the beads while in the magnetic stand. Incubate at room temperature for 30 seconds, and then carefully remove and discard the supernatant. Be careful not to disturb the beads that contain DNA targets.
7. Repeat Step 6 for a total of two washes. Be sure to remove all visible liquid after the second wash. If necessary, briefly spin the tube/plate, place back on the magnet and remove all traces of ethanol with a P10 pipette tip.
8. Air dry the beads for 5 minutes while the PCR tubes are on the magnetic stand with the lid open.
9. Add specified volume of specified buffer and vortex.
10. Incubate samples on the bench for 5 minutes at room temperature.
11. Place samples on a PCR tube (200 μL) DynaMag block.
12. Transfer eluate, now containing the DNA targets, to a new PCR tube using a pipette. Be careful not to disturb the beads.

#### 3.3.4. rSAP (Phosphatase) treatment

1. Remove sample from thermocycler and add 46.75 μL 10 mM Tris HCl pH 8.2 to double the volume to 93.5 μL.
2. Perform an AMPure® XP bead clean up adding 78 μL beads and eluting with 85 μL DD water into a clean PCR tube (see section 3.3.3).
3. Add 10 μL 10 × rCutsmart buffer, mix, then add 5 μL rSAP enzyme.
4. Incubate for 30 min at 37°C.
5. Incubate for 15 min at 65°C.

#### 3.3.5. Removal of 5’-linked Spo11 peptide using recombinant TDP2

1. Perform an AMPure® XP bead clean up adding 83 μL beads and eluting with 50 μL 10 mM Tris HCl pH 8.2 into a clean PCR tube (see section 3.3.3).
2. Add 50 μL 2 × Tdp2 buffer and vortex for 5 seconds.
3. Add 3 μL 30 μM 14M_zTDP2CAT (see section 3.1.3) and vortex for 5 seconds.
4. Incubate at 37°C for 1 hour.

#### 3.3.6. Read 1 adapter ligation

1. During the 60 minute incubation prepare a mastermix of the following components: 15 μL NEBNext® Ultra II Ligation Master, 0.5 μL NEBNext® Ligation Enhancer and keep on ice (see note 11).
2. In a separate tube, mix 1 μL of **Read 1 P5** adapter (15 μM) with 9 μL of adapter dilution buffer and mix thoroughly. This ten-fold diluted stock is sufficient for 8 libraries. Do not add adapters to the master mix.
3. Remove sample from thermocycler and cool to room temperature.
4. Perform an AMPure® XP bead clean up adding 85.5 μL beads and eluting with 26.5 μL 10 mM Tris HCl pH 8.2 into a clean PCR tube (see section 3.3.3).
5. Add 3.5 μL NEBNext® Ultra II End Prep Reaction Buffer.
6. Add 15.5 μL of the NEBNext® master mix.
7. Add 1.25 μL of diluted Read 1 P5 adapter (1.5 μM).
8. Mix thoroughly by vortexing and quickly spin down.
9. Incubate at 20°C overnight in a thermocycler with the heated lid off.

#### 3.3.7. PCR Amplification

1. Perform an AMPure® XP bead clean up adding 78 μL beads and eluting with 15 μL 10 mM Tris HCl pH 8.2 into a clean PCR tube.
2. To 15 μL elution, add 25 μL NEBNext® Ultra II Q5 Master Mix, 5 μL Index Primer (use a unique index primer for each sample) and 5 μL Universal PCR Primer. Vortex for 2 seconds to mix.
3. Perform a PCR reaction with the following steps:
4. After the PCR reaction add 50 μL 10 mM Tris HCl pH 8.2.
5. Make up 1 mL of 0.1 × TE pH 7.6 by mixing 100 μL 1 × TE pH 7.6 with 900 μL DD water.
6. Perform an AMPure® XP bead clean up adding 83 μL beads and eluting with 30 μL 0.1 × TE pH 7.6 into a clean PCR tube (see section 3.3.3).

| Temperature | Time | Cycles |
| --- | --- | --- |
| 98°C | 30 sec | 1 |
| 98°C<br>65°C | 10 sec<br>75 sec | 15 |
| 65°C | 5 min | 1 |
| 4°C | Hold |  |

#### 3.3.8. Library quantification and quality control

1. Prepare a high-sensitivity Bioanalyzer 2100 DNA chip following manufacturer’s protocol
2. Pipette 1 μL of each library into a clean tube containing 9 μL DD water and mix thoroughly (1 in 10 dilution).
3. Carefully load 1 μL of each diluted library onto a Bioanalyzer chip.
4. Load and run the chip on the Bioanalyzer machine. A typical Bioanalyzer 2100 trace is shown in Figure 4 (see note 12).

**Figure 4.**
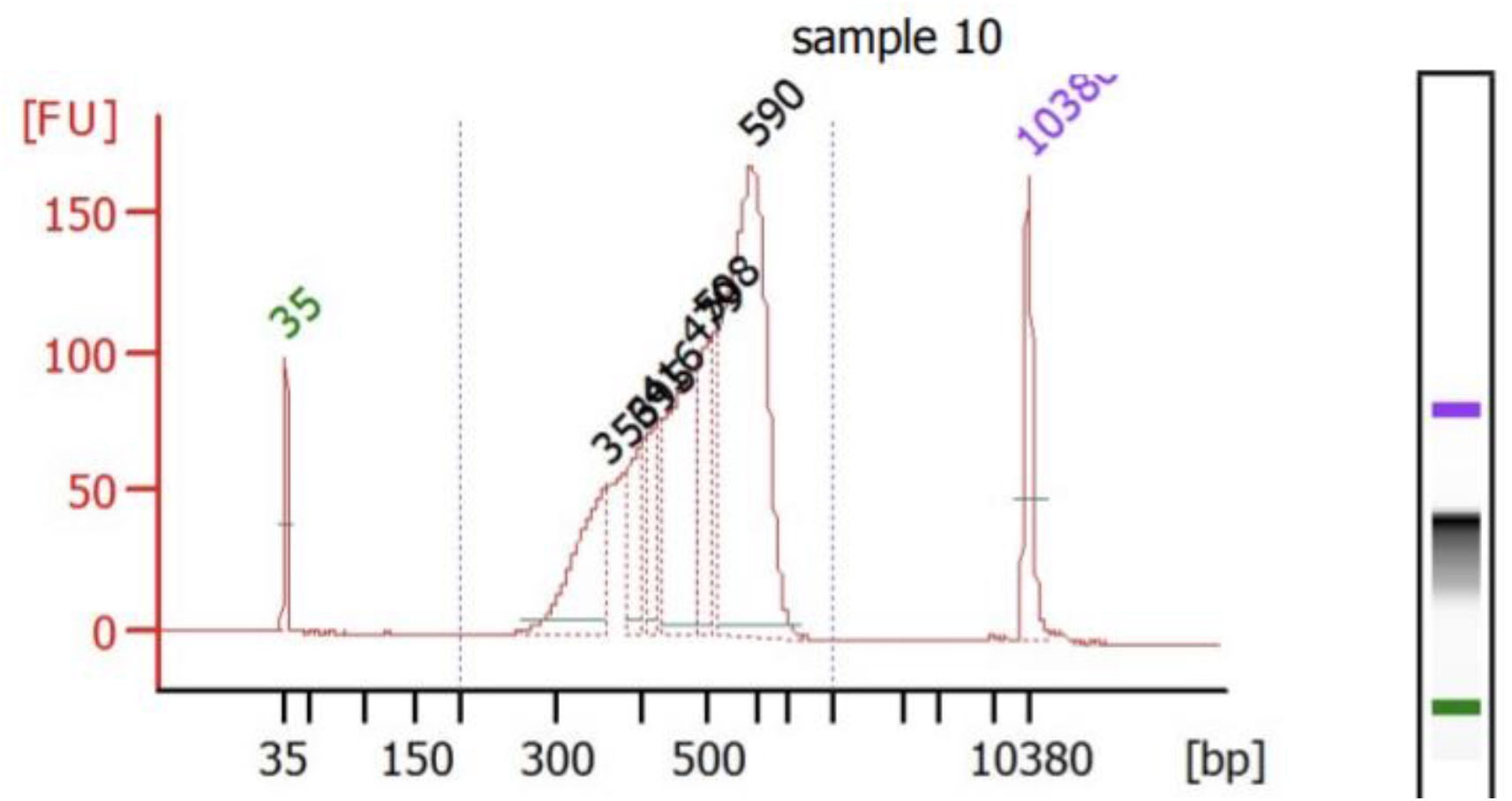
A typical CC-seq library bioanalyzer 2100 trace. Samples were diluted 1 in 10 after PCR and AMPure® XP bead clean up.

#### 3.3.9. Library Sequencing

1. Dilute library in DD water as recommended by the NGS manufacturer.
2. Libraries are compatible with any Illumina®-compatible DNA sequencer (e.g. MiSeq^TM^, NextSeq^TM^ or HiSeq^TM^. Multiple libraries (each with unique indexes, added at the PCR stage) can be multiplexed together (see note 13).

### 3.4 NGS data processing

1. Download the demultiplexed FASTQ files
2. Align the FASTQ files to the genome of interest, for example “sacCer3”, using terminalMapper, https://github.com/Neale-Lab/terminalMapper, a data processing pipeline that aligns via Bowtie2, then calls the read 1 (Spo11-CC coordinate) for each molecule, generating “FullMap” files.
3. Use R or other analysis scripts to plot and analyse the data to explore where Spo11-CCs form (e.g. using functions within the CCTools package, https://github.com/Neale-Lab/CCTools, see Figure 2).

## 4. Notes

1. We recommend that at least two biological replicates (independent cultures) should be prepared per genotype being analysed. Here, we show that CC-seq biological replicate samples have high reproducibility (Pearson’s r = 0.99), have high sensitivity and specificity for Spo11 at nucleotide resolution Figure 2A, 2B, 2C).
2. We have found, empirically, that DNA concentration affects sonication efficiency. Therefore, sonication timing or intensities might have to be adjusted depending on DNA concentration. It is important to aim for all DNA fragments to be below 1 kb, and the majority in the range between 200 and 700 bp.
3. To maintain EtBr staining in the gel use a running buffer containing 1 × TAE with 0.5 μg/mL EtBr. We have also, empirically, found that the concentration of EtBr matters and that low concentrations of EtBr effects with DNA running speeds. We believe that the RNase binds to the DNA and impedes movement of DNA through agarose and that high concentrations of EtBr reduce ionic interactions between proteins and DNA.
4. Excess human CC-enriched DNA can be stored at -80°C for future preparation of additional human:lambda spike material.
5. The final concentration of the spike-in can in theory be lower but must be greater than 50 μg/mL in order to spike in libraries up to a DNA concentration of 100 μg/mL.
6. Final OD_600_ value of ∼6.5 is usually very robust. However, if OD_600_ values differ significantly between cultures processed on the same day sample volumes can be normalised with DD water to balance the centrifuge.
7. DNA samples can be stored in the freezer (-20°C) for up to 2 weeks.
8. DNA concentrations should be between 15 μg/mL and 100 μg/mL
9. DNA concentrations should be between 15 ng/mL and 100 ng/mL
10. It is important to not over dry the AMPure® XP beads because this affects DNA recovery. Bead drying times may have to be adjusted based on the number of samples or pipetting time. The beads should appear “glossy” after the drying stage, if the beads appear “cracked” then they have been overdried.
11. If required, the NEBNext® Master Mix can be made up to 8 hours before use.
12. Library concentrations obtained should be between 20 and 200 nM.
13. Aim for at least 2 million reads per library. Although most library to library differences can be detected with much less reads ∼ 0.5 million.

